# The olfactory bulb reflects structural plasticity within a genetically stable olfactory network

**DOI:** 10.1101/2025.08.08.668515

**Authors:** Tora Olsson, Akshita Joshi, Martin Schaefer, Artin Arshamian, Thomas Hummel, Johan Lundström, Fahimeh Darki

**Author notes:** Corresponding author: Fahimeh Darki, PhD, Department of Clinical Neuroscience, Karolinska Institutet, Nobels väg 9, 17177 Stockholm, Sweden.

## Abstract

The olfactory bulb (OB), the first central relay of the olfactory pathway, plays a critical role in odor perception and exhibits remarkable structural plasticity shaped by environmental influences. This raises a fundamental question about the extent to which the structure of the OB is genetically determined. While both OB volume and function have been implicated in a range of neurodegenerative and neuropsychiatric disorders, such as Parkinson’s disease, schizophrenia, and depression, all of which involve known genetic risk factors, the heritability of the OB structure remains poorly understood. Here, we investigated the heritability of OB volume and the broader olfactory network architecture in a large sample of healthy young adults (n = 941; aged 22-35 years), including monozygotic and dizygotic twin pairs. Using a deep learning-based segmentation model, we first automatically segmented OB volumes and then employed a support vector machine framework to classify zygosity based on within-pair morphological similarity. The OB volume alone showed weak classification performance in distinguishing between monozygotic and dizygotic twins, suggesting limited heritability. However, integrating OB volume with morphometric features from olfactory-associated brain regions, including the hippocampus, parahippocampal gyrus, entorhinal cortex, and medial orbitofrontal cortices, substantially improved classification performance. This effect was specific to the olfactory network, underscoring the distributed nature of genetic influence in this system. These findings suggest that the OB, despite being a highly plastic and environmentally responsive structure, is embedded within a genetically coordinated neural network.

## INTRODUCTION

The olfactory bulb (OB) is a critical structure in the human brain, serving as the first relay station for olfactory information transmitted from peripheral sensory neurons to higher-order brain regions ^1–4^. Beyond its fundamental role in odor perception, alterations in OB structure and function have been increasingly associated with various neurological and psychiatric disorders, including Parkinson’s disease ^3,5^, schizophrenia ^6^, and depression ^7^ – conditions that are known to have strong genetic components. While several genetic studies have identified genes involved in OB development ^8,9^, our understanding of the heritability of OB volume in the general population remains limited.

The human OB exhibits a remarkable degree of plasticity throughout adulthood, capable of structural and functional reorganization in response to illness, sensory experience, training, and deprivation ^10^. This plasticity is demonstrated on a macroscopic level through several key findings. First, OB volume is significantly decreased in patients with a wide range of olfactory disorders, including post-traumatic, post-infectious, sinonasal, and idiopathic olfactory loss ^11–19^. Second, this volume reduction is recoverable, increasing with olfactory training and following the resolution of olfactory disorders ^20,21^. Compellingly, the recovery of olfactory function is associated with a corresponding increase in OB volume, providing strong evidence of its capacity for functionally relevant structural change ^10,22^.

This requirement for plasticity probably arises from the organization of the peripheral olfactory system. Olfactory receptor neurons (ORNs) are uniquely positioned, extending into the nasal cavity where they are directly exposed to the external environment, making them particularly vulnerable to toxins, pathogens, and airborne pollutants. This exposure results in ongoing damage and neuronal turnover ^10^. To maintain function, the olfactory neuroepithelium continuously regenerates ORNs throughout life ^23^. These newly generated neurons must project axons to the olfactory bulb and establish precise synaptic connections, necessitating a high degree of structural and functional plasticity in the bulb to accommodate and integrate these dynamic inputs. Critically, this plasticity is thought to be partially regulated by these centripetal influences, proposing that the OB structural integrity is dependent on this sensory input from the olfactory epithelium ^10^.

While these environmental and experiential factors are well-characterized, the degree to which OB volume is genetically determined remains poorly understood. Heritability estimates of brain structures have played an essential role in identifying genetic contributions to neuroanatomy and informing models of brain development and disease vulnerability ^24,25^.

While extensive research has quantified the heritability of a variety of brain volumes (e.g., the hippocampus and amygdala) ^26^, the OB has remained largely understudied in this context. Consequently, it is unclear whether OB volume reflects primarily genetic variation, environmental influences, or an interaction of both, underscoring the need to disentangle these contributions.

Importantly, olfactory processing is not confined to the OB alone. Instead, it relies on a distributed network of anatomically and functionally connected brain regions, including hippocampus, amygdala, parahippocampus, insula, entorhinal, and orbitofrontal cortex, that collectively support olfactory function ^27–29^. However, whether this broader olfactory network exhibits a coherent pattern of genetic covariation has yet to be explored.

In this study, we address two key questions: (1) is OB volume heritable in a general population of young adults, and (2) does heritability extend to the broader olfactory network, suggesting a genetically coordinated system. A major methodological barrier to addressing these questions has been the challenge of accurately and automatically segmenting the OB. To overcome this limitation, we first validated a deep learning-based segmentation model for automated OB delineation ^30^ using two independent datasets with manually segmented OB volumes. We then applied the automated segmentation pipeline to the Human Connectome Project (HCP) dataset ^31^, a large sample of healthy young adults (n = 1101), including monozygotic (MZ) and dizygotic (DZ) twins, non-twin siblings, and unrelated individuals.

To quantify heritability, we employed a support vector machine (SVM) classification approach based on within-pair morphological similarity. This method, previously applied in neuroimaging studies to detect heritable traits ^32^, is based on the principle that heritable structures should show greater similarity among MZ twins than among DZ twins, non-twin siblings, or unrelated individuals. We applied this framework to assess the heritability of OB volume alone, and in combination with olfactory-associated brain regions. Additionally, we performed similar analyses for other sensory systems by combining OB volume with structural features from the visual and auditory networks to evaluate the specificity of the heritability signal to the olfactory system.

Together, this approach enables a comprehensive investigation into whether the olfactory system exhibits a unique pattern of structural heritability and whether the OB’s heritable signal is amplified when considered within the broader olfactory-associated network.

## METHODS

### Segmentation of the OB

The OB segmentation was performed using an automatic deep learning–based method developed by Desser et al. ^30^. This approach employs a 3D U-Net convolutional neural network trained to segment the OB from combined T1- and T2-weighted MRI data. The model enables fully automated, high-resolution OB segmentation. Full details of the segmentation pipeline and training procedures can be found in the original publication ^30^.

### Validation of Deep Learning-Based OB Segmentation

To evaluate the performance of the deep learning-based segmentation of the OB, we validated the method using two independent datasets. The first dataset included 47 subjects with T1- weighted axial whole-brain scans and high-resolution T2-weighted coronal images. MRI data were acquired on a 3T Siemens Verio scanner (Erlangen, Germany). The T1-weighted MPRAGE sequence had the following parameters: TR = 2300 ms, TE = 2.98 ms, flip angle = 9°, field of view (FOV) = 240 × 256 mm^2^, acquisition matrix = 250 × 256, voxel size = 1 × 1 × 1 mm^3^, and 176 slices. The T2-weighted high-resolution coronal sequence was acquired with TR = 5500 ms, TE = 110 ms, flip angle = 150°, FOV = 120 × 120 mm^2^, matrix = 256 × 256, voxel size = 0.47 × 0.47 × 1.2 mm^3^, and slice thickness = 1.2 mm. The second dataset comprised 33 participants. Data were collected using a 3T Siemens Prisma scanner (Erlangen, Germany) with a 32-channel head coil. T1-weighted 3D MPRAGE scans were acquired using the following parameters: TR = 2300 ms, TE = 3.43 ms, flip angle = 9°, voxel size = 1 × 1 × 1 mm^3^, 160 slices, and GRAPPA acceleration (factor = 2). The T2-weighted turbo spin echo sequence was acquired with TR = 6200 ms, TE = 117 ms, flip angle = 160°, voxel size = 0.2 × 0.2 × 1 mm^3^.

In both datasets, the left and right OB were initially segmented manually by two expert raters. Segmentations differing by more than 10% were re-segmented by a third rater. Finally, the manually segmented OB volumes were averaged to obtain the final volume estimates. These datasets provided a robust basis for validating the deep learning-based OB segmentation method across different scanners and acquisition protocols. We assessed the model’s generalizability by computing correlations between the automated and manual OB volume measurements.

### Deep Learning-Based OB Segmentation of the Human Connectome Project (HCP) Data

To determine the OB volume of healthy individuals and investigate the heritability of the OB volume, we utilized structural MRI data from the Young Adult HCP dataset, which includes high-resolution T1- and T2-weighted scans from approximately 1,200 healthy participants aged 22 to 35 years ^31^. This cohort is particularly suitable for heritability analyses as it includes MZ and DZ twins, non-twin siblings, and unrelated individuals, enabling well-powered assessments of genetic and familial contributions to neuroanatomical variations.

Automated OB segmentation was performed on 1101 subjects with both T1- and T2-weighted images in native subject space. To ensure the anatomical plausibility and reliability of OB segmentations, we implemented a rigorous post-processing quality control procedure to identify and exclude erroneous results. First, participants were excluded if the voxel distance (*D)* between the left and right segments was either *D* ≤ 0 or *D* ≥ 8, indicating no separation or unrealistically large separation between the left and right bulbs. Second, participants were discarded if both OBs were located within the same hemisphere. Third, participants were excluded if one bulb had volume <= 15 mm^3^ and the absolute volume difference between the left and right OB volume exceeded 10 mm^3^, suggesting segmentation failure on one side. Applying these criteria resulted in the exclusion of 160 participants, leaving a total of 941 segmented OBs in the final dataset. All pre-processing, segmentation, and post-processing steps were performed using Python version 3.7.

To characterize the spatial distribution and anatomical variability of OBs in the healthy population of young adults, we also generated a probabilistic OB map by spatially normalizing the segmentations and averaging the binary OB masks across all included participants. This map provides a population-level reference for OB location and extent, supporting further analyses.

### Support Vector Machine (SVM) Classification for Heritability Assessment of OB

To assess the heritability of OB volume, we employed a SVM classification approach similar to that used by Miranda-Dominguez et al. ^32^. This method is based on the principle that heritable traits manifest as greater similarity among individuals with a higher degree of genetic relatedness. Therefore, by comparing the similarity of brain features between MZ twins, DZ twins, non-twin siblings, and unrelated individuals, we can infer the extent to which genetic factors influence the OB volume.

To account for individual differences in overall brain size, the left and right OB volumes were first adjusted for intracranial volume (ICV) using linear regression. The resulting standardized residuals were then used in all subsequent analyses. The dataset was divided into four groups: 208 MZ twins forming 104 MZ pairs, 116 DZ twins forming 58 DZ pairs, 212 non-twin siblings forming 106 sibling pairs (excluding families with twins to prevent group overlap), and 128 unrelated individuals forming 64 unrelated pairs (selected to have no siblings in the dataset). For each twin and sibling pair, we calculated the absolute difference in standardized OB values between the two individuals. For the unrelated group, we created age-matched scan pairs by randomly selecting unrelated individuals without familial overlap and computing the absolute differences between each pair. These pairwise difference values served as input features to train SVM classifiers to distinguish between different types of genetic relationships (e.g., MZ vs. DZ) with an 80/20 train/test split. The assumption is that more heritable traits will result in smaller differences between MZ twins relative to DZ twins. Model performance was assessed using five-fold cross-validation, and permutation testing (5,000 permutations) was applied to evaluate statistical significance.

### Heritability Analysis of Olfactory System Structures

Given that the olfactory system functions as a developmentally and functionally integrated network, we hypothesized that the involved brain regions are collectively shaped by genetic influences. Thus, we extended the SVM classification framework to combine OB with brain regions structurally and functionally associated with olfactory processing, specifically: bilateral hippocampus and amygdala, parahippocampal gyri, medial orbitofrontal, insula, and entorhinal cortices ^28,29^. Regional measures for these structures were extracted from the HCP Freesurfer outputs, which provide standardized, high-resolution cortical and subcortical segmentations ^31,33^. Although the piriform cortex is a key region in primary olfactory processing, it was not included in the analysis because it is not represented in standard Freesurfer parcellation schemes.

Bilateral OB volumes, along with 12 additional features, were entered into the same SVM classification framework (5-fold cross-validation) for classifying MZ twins from DZ twin pairs. These 14 features yielded 2^14^ − 1 = 16,383 unique feature combinations, ranging from single-feature models to those including all features. This approach allowed us to systematically evaluate the collective heritability of the broader olfactory network by identifying combinations of features that best distinguished genetically identical from non-identical twin pairs.

### Control Analyses Using Visual and Auditory Networks

To evaluate whether the heritability of the olfactory system reflects a system-specific pattern rather than a general effect of other brain regions, we performed control analyses using unrelated sensory networks. We hypothesized that combining OB volume with regions from unrelated sensory systems, such as visual or auditory networks, would yield lower classification performance, reflecting weaker shared developmental and genetic integration with the OB.

For the visual network, we included structural measures from regions known to support visual processing: the cuneus, fusiform gyrus, lateral occipital cortex, lingual gyrus, pericalcarine cortex, and precuneus. For the auditory network, features included the inferior temporal gyrus, middle temporal gyrus, pars triangularis, superior temporal gyrus, supramarginal gyrus, and transverse temporal gyrus. All regional measures were extracted from the HCP Freesurfer outputs ^31,33^. In both control analyses, the left and right OB volumes were included and combined with these sensory-specific features. Each feature set (OB + visual regions, OB + auditory regions) was analyzed using the same SVM classification framework (5-fold cross-validation) used in the olfactory network analysis. A total of 14 features were considered in each control analysis, resulting in 16,383 possible feature combinations per network.

## RESULTS

### Validation of Deep Learning Segmentation of the Olfactory Bulb

We applied a deep learning-based segmentation pipeline ^30^ to automatically segment the OB in two independent datasets. Each dataset also included manual segmentations performed by expert raters, providing reference volume measurements for comparison.

To evaluate the model’s performance, we computed Pearson correlations between the automated and manual volumes of the left and right OBs. As shown in Fig. 1A, the correlation coefficients for each dataset separately illustrate the consistency of model performance across the two independent datasets. We then combined the two datasets and plotted scatter plots comparing manual and automated segmentations for the left and right OBs (Fig. 1B). The results demonstrated agreement between manual and deep learning-based segmentation volumes. In the combined dataset, the Pearson correlation coefficient was *r* = .76 (*p* = 2.2 × 10^−14^) for the left OB and *r* = .70 (*p* = 1.7 × 10^−11^) for the right OB, indicating reliable performance of the deep learning–based segmentation.

**Figure 1.**
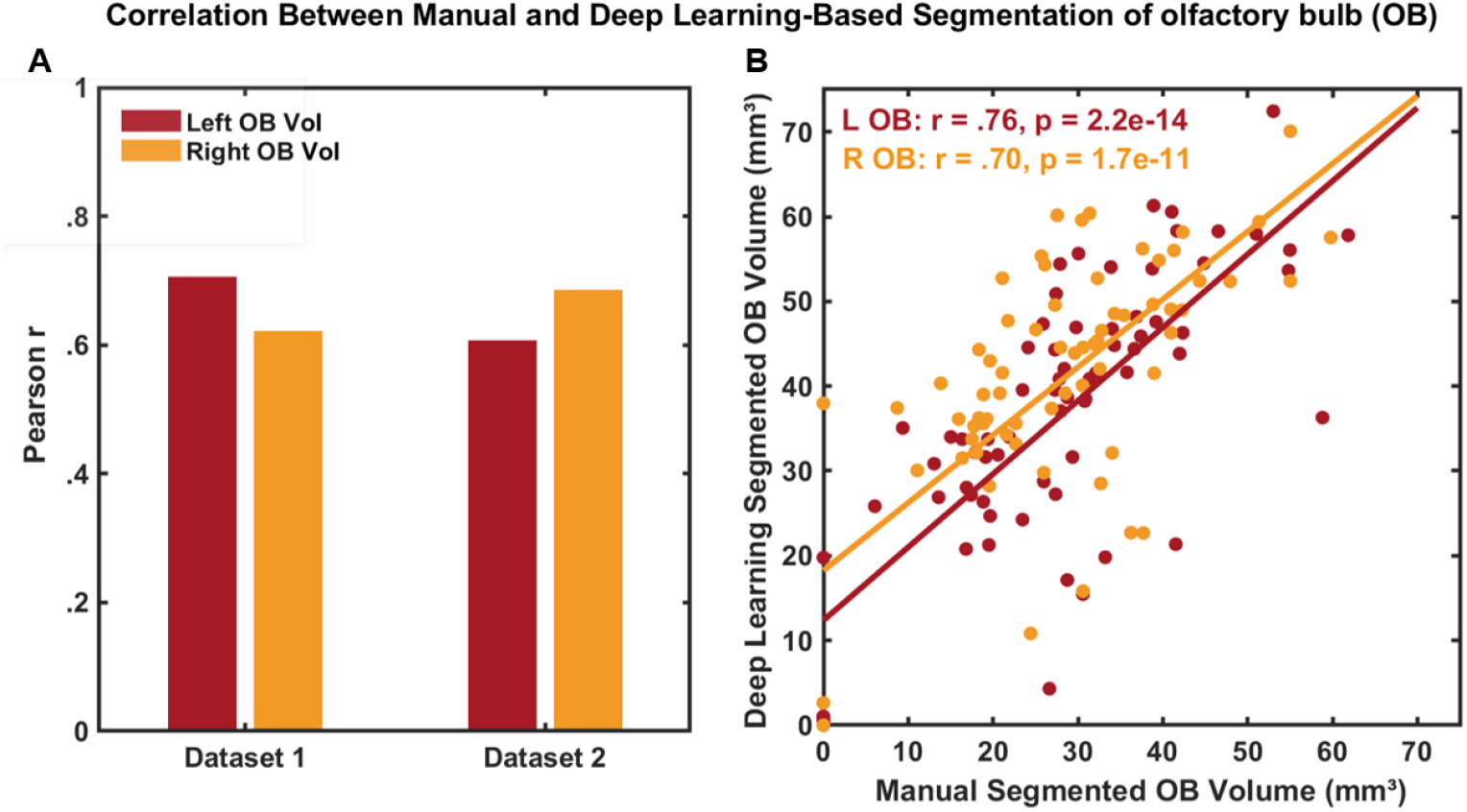
Validation of Deep Learning-Based Olfactory Bulb (OB) Segmentation Against Manual Annotations. (A) Bar plot showing Pearson correlation coefficients between manual and automated OB volume segmentations for left and right OB, separately in two validation datasets. (B) Combined scatter plot illustrating the correlation between manual and deep learning–based OB volumes for the left and right OB across both datasets. Each dot represents a single subject.

### Olfactory Bulb Segmentation Across the HCP Cohort

After demonstrating the reliability of the deep learning segmentation model, we next applied the segmentation model to a large sample of healthy young adults from HCP. This dataset includes genetically informative participants (e.g., twins, siblings, and unrelated individuals), making it well-suited for investigating the heritability of the OB.

Automated OB segmentation was successfully performed across 941 HCP subjects. Using the mask of segmented OB, we generated a probabilistic OB map in standard space, illustrating the voxel-wise likelihood of OB presence across individuals (Fig. 2A). The resulting map demonstrated high spatial consistency and anatomical plausibility, confirming the model’s robustness in a healthy, high-resolution population sample. These segmentations in a large-scale sample provided a basis for subsequent heritability analyses, enabling the investigation of genetic contributions to variation in OB volume within the HCP twin cohort.

**Figure 2.**
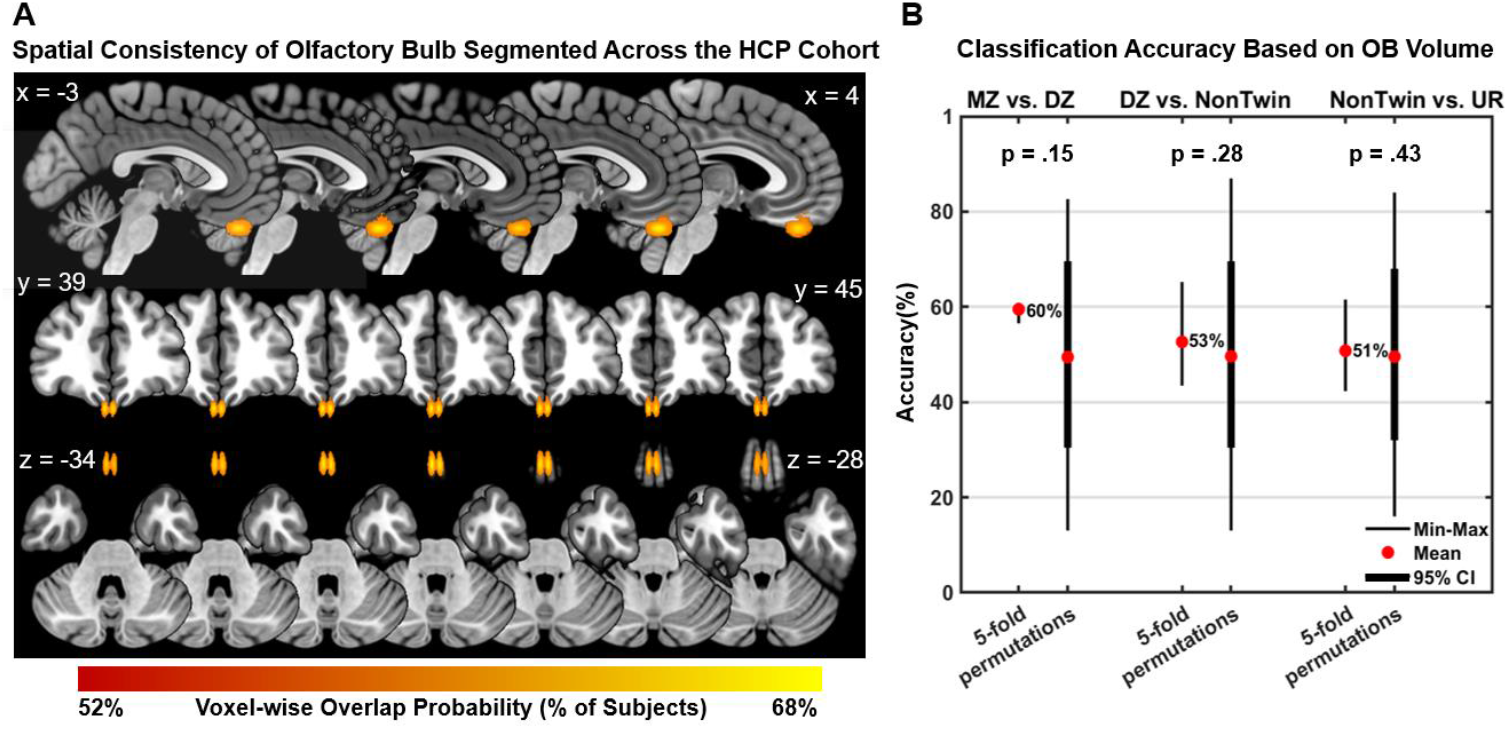
Spatial consistency of olfactory bulb (OB) segmentation and classification of genetic relatedness of OB volume. (A) Probabilistic map of OB segmentations in the Human Connectome Project (HCP) dataset. The map represents the voxel-wise probability of OB presence, defined as the proportion of subjects in which each voxel was labeled as OB following deep learning-based segmentation. Warmer colors indicate regions with higher spatial overlap, reflecting consistent OB localization across individuals. (B) Classification accuracy across groups using OB volume. Bar plots represent 5-fold cross-validated classification accuracy and permutation-based null distributions for three pairwise classifications: monozygotic (MZ) vs. dizygotic (DZ) twins, DZ twins vs. non-twin siblings, and non-twin siblings vs. unrelated (UR) individuals. Only the left and right OB volumes were used as features in these models. Thin lines indicate the full range of observed accuracy, while thick lines represent 95% confidence intervals (CI).

### Moderate Accuracy for OB Volume Differentiation Between MZ and DZ Twins

We next examined whether bilateral OB volume reflects genetic relatedness by analyzing the bilateral OB volumes. A SVM classifier with 5-fold cross-validation was used to distinguish between different levels of genetic relatedness: MZ vs. DZ twins, DZ twins vs. non-twin siblings, and non-twin siblings vs. unrelated individuals. For each pair of individuals within each group, the absolute difference in standardized OB volume measures was computed and used as input to the classifier. The classification between MZ and DZ twins yielded an accuracy of 60%, suggesting a low genetic contribution to OB volume, although the result did not reach statistical significance (*p* = .15, permutation test; Fig. 2B). Classifications between DZ twins and non-twins, and between non-twins and unrelated individuals, showed chance-level accuracies (∼ 50%), indicating minimal volumetric differences in these groups.

### Multiregional Olfactory Network Features Improve Classification of Genetic Relatedness in Twins

Given the observation that OB volume showed none or at best low heritability alone, we next extended our analysis to investigate whether combining OB volume with structural measures from other olfactory-related regions would yield a more robust signal, reflecting the olfactory system’s role as a developmentally linked and genetically regulated network. Structural measures from both the left and right hemispheres of the regions involved in olfactory processing were included as features: OB, hippocampus, amygdala, parahippocampal gyrus, insula, entorhinal cortex, and medial orbitofrontal cortex, resulting in a total of 14 features. To fully explore regional contributions, we performed classification across all possible feature combinations (n = 16,383).

This analysis identified 76 feature combinations that achieved classification accuracy greater than 75%, with all reaching statistical significance (*p* < .006, permutation test with 5,000 iterations; Fig. 3A). Of these, 67 combinations included either the left, right, or bilateral OB, highlighting its central role in distinguishing between MZ and DZ. These results indicate that integrating OB volume with structural measures from key olfactory network regions significantly enhances the heritable signal. Among the high-performing models, the combination of OB volume with bilateral hippocampus volume, right parahippocampal gyrus, right medial orbitofrontal cortex, and left entorhinal cortex thickness was the top-performing, achieving the highest classification accuracy (82%, *p* = .004; Fig. 3A). Importantly, this combination did not yield significant classification accuracy when distinguishing DZ twins from non-twin siblings, nor when classifying non-twins versus unrelated individuals (Fig. 3B), indicating that the observed effects are specific to the genetic relatedness captured by the MZ vs. DZ comparison.

**Figure 3.**
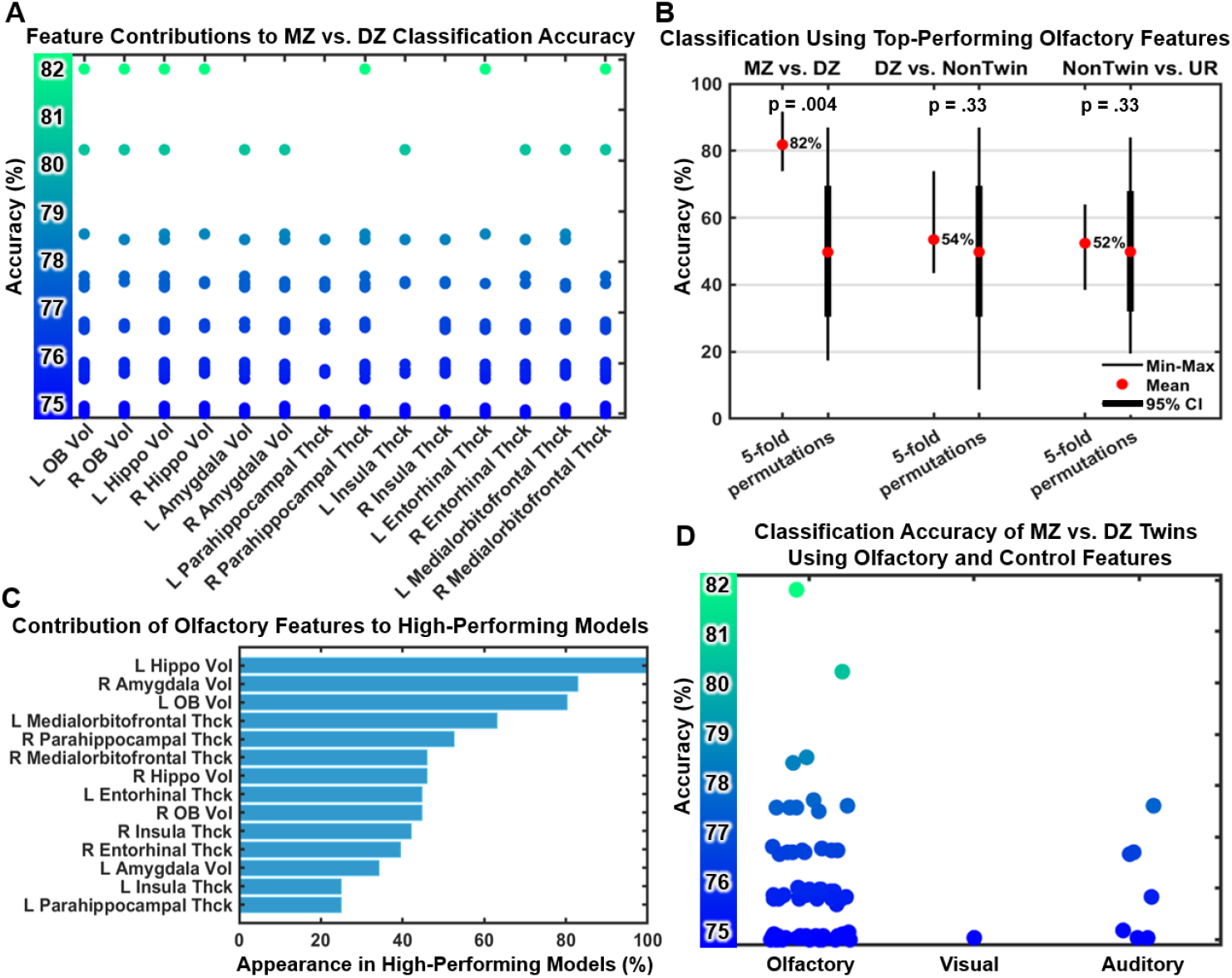
Classification performance and feature contribution. (A) Scatter plot visualizing classification accuracies and feature contribution for distinguishing monozygotic (MZ) vs. dizygotic (DZ) twins. Each dot represents a feature contributed to one of 76 high-performing classifications (accuracy > 75%). The y-axis shows the classification accuracy per feature (ranging from 75% to 82%), and the x-axis lists the olfactory features. The color of each dot reflects the accuracy of the feature combination, lighter shades indicating higher performance. (B) Classification accuracy across groups using top-performing feature combination (i.e., bilateral OB, bilateral hippocampus volume, right parahippocampal gyrus, right medial orbitofrontal cortex, and left entorhinal cortex thickness contributed to an accuracy of 82% for distinguishing MZ vs. DZ). Bar plots show 5-fold cross-validated classification accuracies and permutation-based null distributions for three group comparisons: MZ vs. DZ twins, DZ twins vs. non-twin siblings, and non-twin siblings vs. unrelated individuals (UR). Thin lines indicate the full range (minimum to maximum) of observed accuracy, while thick lines represent 95% confidence intervals (CI). (C) Percentage of appearance for each olfactory feature across high-performing models. Barplot illustrates the percentage of times each feature appeared among the 76 feature combinations that achieved classification accuracy >75%. (D) Comparison of classification accuracy for distinguishing MZ vs. DZ twins using olfactory, visual, and auditory features combined with OB volume. Jittered colored dots represent individual model performances for all feature combinations that achieved accuracy above 75% (76 olfactory feature combinations, one visual feature combination, and 7 auditory feature combinations). The colorbar indicates classification accuracy.

Beyond the OB, left hippocampus, right amygdala, and left medial orbitofrontal cortex contributed most consistently to high-performing classification models (accuracy >75%; Fig. 3C). Their frequent involvement across high-performing models supports the hypothesis that heritability in the olfactory system is not localized to the OB alone but rather emerges from a distributed network with genetically coordinated development.

### Limited Heritability Signal from OB Combined with Non-Olfactory Sensory Regions

Based on our hypothesis that the olfactory system is collectively heritable, we predicted that combining OB volume with regions from other sensory systems would not yield comparable classification performance. To test this, we conducted control analyses using structural measures from the visual and auditory networks. The left and right OB volumes were separately combined with structural measures of the visual (e.g., cuneus, fusiform, lateral occipital, lingual, pericalcarine, and precuneus) and the auditory (e.g., inferior temporal, middle temporal, parstriangularis, superior temporal, supramarginal, and transverse temporal) networks. These features were separately entered into the same SVM classification framework (5-fold cross-validation) for classifying between MZ and DZ. For each network, a total of 14 features were considered, resulting in 16,383 possible feature combinations per analysis.

In contrast to olfactory network which yielded 76 models with classification accuracies exceeding 75% in distinguishing MZ from DZ twins, control analyses using visual and auditory features resulted in fewer high-performing models (Fig. 3D). For visual network, only one feature combination exceeded the 75% accuracy threshold, involving left and right OB, lingual, precuneus, and left fusiform (*p* = .003, permutation test 5000 permutations). For the auditory network, 7 feature combinations yielded an accuracy above 75% (maximum accuracy = 77%; *p* = .003, permutation test). The features involved in these combinations were less frequent compared to olfactory models, supporting the notion that the heritability of OB volume is specifically tied to its integration within the olfactory system rather than driven by general sensory or structural brain features.

### Visualization of Twin-Pair Similarity Across Olfactory Structures

To further characterize the evidence of heritability in olfactory structures and complement the classification findings, we visualized within-pair similarity across all groups (MZ, DZ, NonTwin, and unrelated individuals) using the features most predictive of distinguishing MZ from DZ twins. Specifically, we focused on the top-performing feature combination that achieved an MZ vs. DZ classification accuracy of 82% (*p* = .004, permutation test). This feature set included bilateral OB volume, left and right hippocampus volume, right parahippocampal gyrus volume, right medial orbitofrontal cortex thickness, and left entorhinal cortex thickness (Fig. 3A). To evaluate the added value of the broader feature set, we directly compared this feature combination with OB volume features alone. Intra-pair similarity was quantified using the mean absolute standardized difference and visualized for both (1) the mean of bilateral OB volume and (2) the mean of features contributing to the top-performing model. While MZ twins consistently exhibited the lowest intra-pair differences in both comparisons, the mean value of the top-performing feature set showed greater separation between MZ and other groups, indicating stronger evidence of heritability (all *p* < 0.001; Fig. 4A). This pattern was further supported by correlation analyses: MZ pairs demonstrated higher within-pair correlations for the top-performing feature set (r = .70) compared to OB volume alone (r = .41), whereas DZ showed weaker associations (Fig. 4B). Together, Fig. 4 supports the findings by illustrating that although OB volume exhibits moderate heritability, the combination OB volume with multiple olfactory-related features reveals a more robust heritability signal.

**Figure 4.**
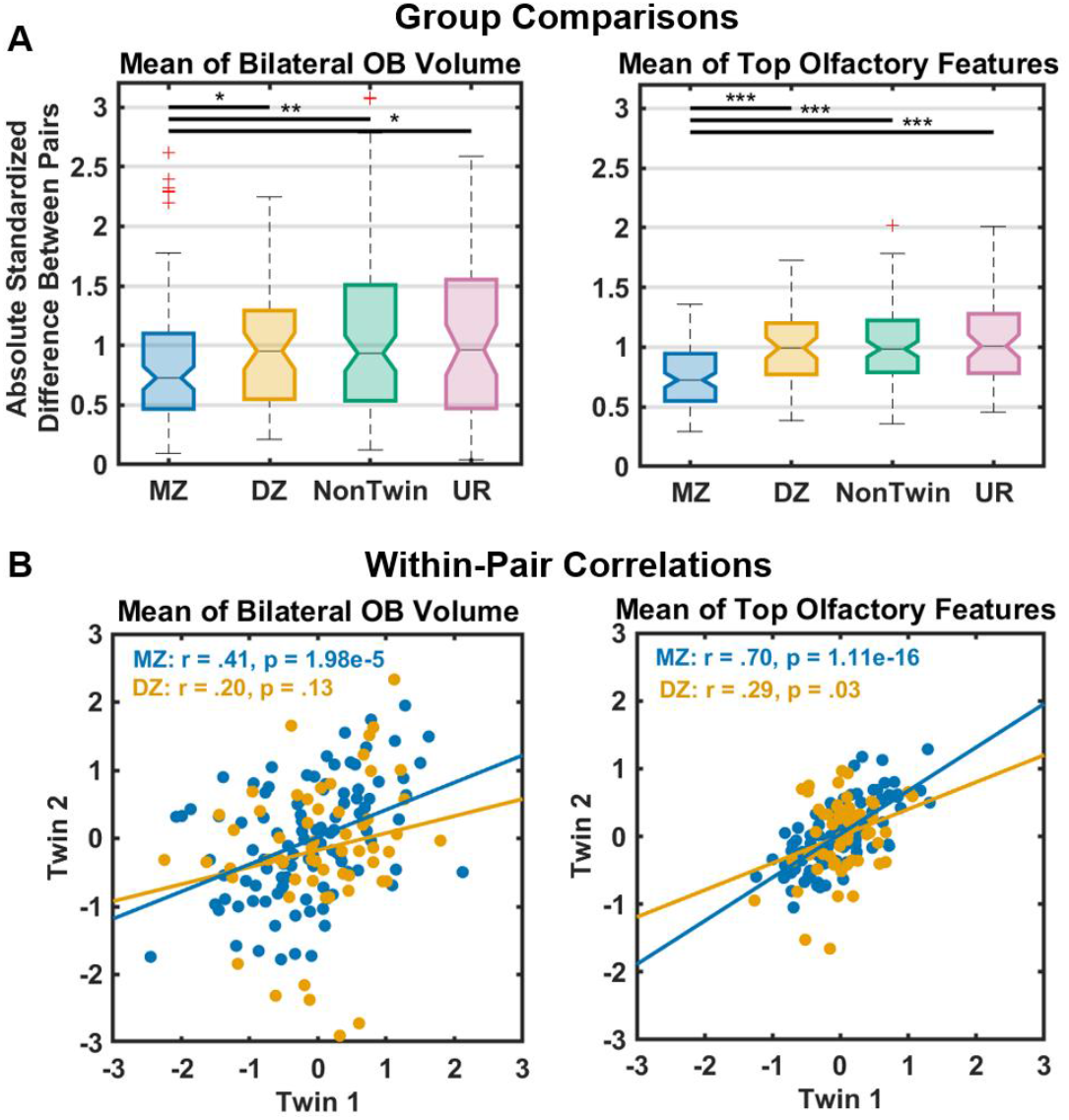
Group comparisons and within-pair correlations for (left) the mean of bilateral olfactory bulb (OB) volume and (right) the mean of the olfactory features contributing to the top-performing model for distinguishing monozygotic (MZ) vs. dizygotic (DZ) twins (accuracy = 82%, features: bilateral OB, bilateral hippocampus volume, right parahippocampal gyrus, right medial orbitofrontal cortex, and left entorhinal cortex thickness). (A) Box plots group-wise distributions of mean absolute standardized differences between pair members for (left) bilateral OB volume and (right) all features contributed to the top-performing classification. Groups include MZ twins, DZ twins, nontwin siblings, and unrelated individuals (UR). Asterisks indicate statistically significant pairwise group differences (* *p* < 0.05, ** *p* < 0.01, *** *p* < 0.001). (B) Scatter plots show within-pair correlations for MZ and DZ twins for (left) the mean of bilateral OB volumes and (right) the mean of all features contributed to the top-performing classification.

## DISCUSSION

In this study, we investigated the heritability of the human OB volume using high-resolution MRI data from the HCP data. We show that the OB volume alone was not sufficiently heritable to enable reliable classification of zygosity (accuracy = 60%). This finding is consistent with previous observations that the OB is a plastic node in the olfactory network susceptible to environmental influences, such as viral exposure, inflammation, or individual differences in olfactory experience ^6,7,10^.

Critically, when OB volume was analyzed in conjunction with olfactory-associated regions, classification performance improved substantially. Specifically, we included cortical thickness and gray matter volume measures extracted from FreeSurfer analysis of the HCP dataset, focusing on regions such as the hippocampus, amygdala, parahippocampal gyrus, entorhinal cortex, insula, and medial orbitofrontal cortex. These areas form part of the canonical olfactory network and are known to contribute to olfactory perception, memory, and emotion ^27^. Unfortunately, we were unable to include the piriform cortex, a key olfactory region, due to limitations in FreeSurfer’s standard parcellation scheme, which does not reliably segment this structure. Including this node may have increased the classification accuracy of the models. Our results suggest that heritability within the olfactory system is distributed across multiple olfactory-associated regions and is more effectively captured when the OB is considered as part of an integrated network rather than in isolation. This is consistent with previous findings that many olfactory-related brain regions, especially in the temporal and frontal lobes, exhibit substantial heritability in cortical volume and thickness ^24,25,34^. Moreover, the improvement in classification performance was specific to the olfactory network. Integrating the OB volume with structural features from other sensory systems (i.e., visual and auditory cortices) did not reach consistency in high accuracy. This pattern supports the hypothesis that the OB combined with other regions within the olfactory system forms a genetically coordinated structural network, distinct in its heritability profile from other sensory areas.

Notably, beyond the OB, the left hippocampus, right amygdala, and left medial orbitofrontal cortex were the most frequently selected features in models achieving classification accuracy above 75%. These structures are not only anatomically connected to the OB but also play essential roles underlying idiosyncratic odor perception that is driven by the orbitofrontal cortex ^35^, as well as odor-related memory (e.g., hippocampus), affective processing (e.g., amygdala), and perceptual integration (e.g., insula) ^35–37^. Their consistent appearance across high-accuracy models suggests that heritability in the olfactory system may be supported by a core set of interconnected regions whose coordinated development reflects underlying genetic architecture.

In addition, the plasticity of the OB adds an important interpretive layer. Previous studies have demonstrated that OB volume can change extensively in response to environmental and sensory experience, including olfactory training or deprivation (Petersen et al. 2024; Yan et al. 2022). This suggests that while OB size reflects genetic factors, it is more malleable than adjacent cortical areas, supporting that OB contributes meaningfully to heritable classification only in conjunction with other olfactory regions.

An important consideration in interpreting these findings is the age range of the HCP sample, consisting of young adults between 22 and 35 years of age. While this cohort is ideal for high-quality neuroimaging and twin-based heritability analyses, it also reflects an age range during which environmental exposures may have already shaped brain structure, particularly in sensory systems like olfaction. The OB is known to exhibit structural plasticity in response to factors such as viral infections, allergies, pollution exposure, and olfactory training or deprivation, all of which accumulate with age and may obscure genetically driven variation ^3,7^. These environmental influences could partially explain the relatively weak heritability signal of OB volume alone observed in our study. It is plausible that stronger heritability would be observed in younger populations, such as infants or children, who have not yet undergone extensive environmental shaping of their olfactory systems. Future studies in pediatric cohorts, ideally with longitudinal designs, will be crucial for disentangling the respective contributions of genetic and environmental influences on OB development across the lifespan.

Together, these findings contribute to a growing understanding of the genetic architecture of the olfactory system and highlight the importance of considering distributed network features in heritability analyses. While OB represents a structurally plastic node influenced by both genetic and environmental factors, its integration with key olfactory-associated brain regions reveals a coherent and genetically influenced network.]

## CONFLICT OF INTEREST

The authors report no conflict of interest.

## ACKNOWLEDGEMENTS

This work was supported by the Swedish Research Council (VR 2021-02700) grant awarded to Professor Johan Lundström. We gratefully acknowledge Professor Thomas Hummel and the Smell & Taste Clinic at the Department of Otorhinolaryngology, University of Dresden Medical School, for generously providing the MRI data for validation. For the analysis, data were provided by the Human Connectome Project, WU-Minn Consortium (Principal Investigators: David Van Essen and Kamil Ugurbil; 1U54MH091657) funded by the 16 NIH Institutes and Centers that support the NIH Blueprint for Neuroscience Research; and by the McDonnell Center for Systems Neuroscience at Washington University.

## Notes

### Competing Interest Statement

The authors have declared no competing interest.

